# Decoupled ionic and particulate clearance pathways determine the *in vivo* fate of a synthetic nanoclay-BMP-2 biomaterial during ectopic bone induction

**DOI:** 10.64898/2026.05.01.722265

**Authors:** Yang-Hee Kim, Juan Aviles Milan, Janos Kanczler, Agnieszka A. Janeczek, Richard O.C. Oreffo, Jonathan I. Dawson

## Abstract

Nanoclay-based biomaterials offer promise for localised growth factor presentation, yet their *in vivo* degradation, clearance, and systemic fate remain poorly defined. Here, we investigate the fate of a synthetic nanoclay-BMP-2 gel during ectopic bone induction using a combination of *in vivo* imaging, histology, and component-resolved elemental analysis. Fluorescent tracking confirmed prolonged localisation of BMP-2 within the nanoclay gel and robust bone induction despite negligible growth-factor release. Inductively coupled plasma mass spectrometry (ICP-MS) revealed divergent clearance kinetics for lithium and silicon, structurally distinct components of the clay crystalline lattice, indicating decoupled ionic and particulate degradation pathways. Early clearance was dominated by cell-mediated fragmentation and the transport of clay particulates, while later stages involved preferential lithium release associated with local clay dissolution as well as integration within newly formed bone. Systemic biodistribution analysis demonstrated rapid, transient lithium release into circulation with renal clearance, contrasted with initial hepatic and then later-phase renal handling of silicon species. Together, these findings define a multiphasic *in vivo* clearance model for nanoclay biomaterials consistent with progressive remodelling, localised BMP-2 activity and, importantly, safe systemic handling. This work provides mechanistic insight into the activity and clearance of nanoclay-based regenerative therapies and establishes the importance of component-resolved tracking for evaluating the biodistribution of degradable inorganic biomaterials.

## 1. Introduction

For over six decades, a plethora of studies have focused on developing hydrogels to control the delivery of drugs and growth factors, such as bone morphogenetic protein-2 (BMP-2) for stimulating regenerative responses [1]. Despite exciting progress in many contexts, the safe and effective delivery of BMP-2 persists as a significant unmet clinical need. This need is driven both by the continued reliance of surgeons on BMP-2 for reliable bone fusion and repair - particularly in aged or metabolically compromised patients, together with the well-documented risks of off-target effects. These risks, which include inflammation, heterotopic bone and osteolysis, can be attributed to the limitations of the delivery vehicles in current clinical use compounded by the high doses thereby needed to achieve efficacy[2, 3].

The concept of slow or sustained release, originally employed in pharmacology for controlling the systemic bioavailability of drugs over time, has represented by far the dominant strategy to maximize the local effects of drugs/growth factors on cell stimulation while minimizing off-site and off-target effects. However slow-release strategies still face the challenge of progressive depletion of the molecule from the target site. Furthermore, while slowed release may reduce some off-target effects, local inflammation and heterotopic bone remain a significant risk due to the potency of BMP-2, even at low concentrations. Biomaterials that stably bind and retain (rather than release) active BMP-2 represent an alternative strategy that is arguably more suited to the goal of local tissue regeneration. The effectiveness of this strategy is however reliant on the ability of the biomaterial, not only to bind and stabilize local concentrations of active growth factor, but also to host cellular ingress into the BMP-2-retaining space and then allow subsequent remodelling and/or clearance of the biomaterial by cells as the new tissue forms in its place.

Certain nanoclays such as the synthetic hectorite LAPONITE^®^ (Na_0.7_^+^[(Si_8_Mg_5.5_Li_0.3_)O_20_(OH)_4_]^−0.7^) are promising materials for localised BMP-2 delivery due both to their gel forming ability and high sorptive capacity for biomolecules. Synthetic hectorites comprise disk-shaped crystals of approximately 25 nm diameter and 1 nm thickness possessing a permanent negative surface charge and an amphoteric rim charge in the form of Mg-OH and Si-OH groups on the particle edge [4]. In water, these particles form delaminated dispersions of individual clay disks which generate, at certain solid and ionic concentrations, arrested colloidal gel states driven by a range of repulsive and attractive interactions between the particles. The thixotropic nature of the nanoclay afford useful rheological properties for injectable delivery and *in situ* gelation, and the very high specific surface area of nanoclay colloids (>300 m^2^ /g solid) provides an excess of adsorption sites for biological agents [5, 6]. Thus, studies have shown that nanoclay gels possess broad spectrum ability to adsorb diverse model proteins, achieve loading concentrations in the order of tens of milligrams per millilitre and exhibit negligible release over extended time frames both *in vitro* [5, 7] and *in vivo* [8]. These properties have been applied beneficially for protein delivery across a range of applications (see [9] for a recent review). Results from our and other groups have demonstrated that nanoclay delivery and localisation of BMP-2 specifically, allows effective significant dose reductions [10-12], attenuation of inflammatory effects [13], and efficient bone fusion and repair either via BMP-2 delivery in nanoclay colloidal gels [13-17], or through nanoclay-BMP-2 functionalization of polymeric hydrogels or porous implants [8, 12, 14-16, 18].

The observation of effective bone induction despite negligible BMP-2 release confers, as noted above, a process of progenitor cell invasion into the BMP-2-retaining implant, followed by its subsequent remodelling as bone tissue forms within the space that the gel had occupied. A recent study by Furuichi et al. [13], for example, evidenced in both the classic ectopic bone model and in a posterolateral spinal fusion model, that clay gels serve as a template within and throughout which first cartilage and then bone forms in response to nanoclay gel-retained BMP-2. This contrasted starkly with the conventional BMP-2 loaded absorbable collagen sponge in which bone formed rapidly, and with minimal cartilage formation, exclusively in a narrow region around the surface. The nanoclay gel implant appeared, at least *via* conventional bone histology and computer tomography analysis, to be completely replaced over time by new bone tissue.

While the process of BMP-2 induced bone formation is well understood, the mechanism by which BMP-2 sequestered within nanoclay gel becomes accessible to cells, is not. Also critical is the fate of nanoclay itself, both in terms of its clearance from the implant site over the course of bone formation and its subsequent biodistribution and/or clearance from the body. The fate of a biomaterial within the body is an important safety consideration as persistence or accumulation can have toxicity implications. Furthermore, the process by which cells invade, colonize and remodel what presents as a relatively dense, non-porous - albeit highly hydrated - material (see, for example, scanning electron micrographs in [19]) is not intuitively obvious. While there is *in vitro* evidence that micro-sized aggregates of nanoclay undergo uptake and degradation within even non-phagocytic cells via the endosome-lysosome axis [20-22], the extent and rate to which the colloidal gel is phagocytosed and undergoes dissolution and/or transportation within the body, by the constitutive cells of the foreign body response, remains largely unknown.

Here, we combine component-resolved elemental tracking with histological analysis to define the *in vivo* clearance, remodelling, and systemic fate of a nanoclay-growth factor composite during ectopic bone formation. Following confirmation of BMP-2 retention via *in vivo* imaging of labelled BMP-2 and bone induction *via* micro computed tomography (µ-CT), we applied inductively coupled plasma mass spectrometry (ICP-MS) to examine the rate and kinetics of nanoclay clearance from the implant site and the nanoclay’s subsequent biodistribution within organs and tissues over the course of 8 weeks. By independently following lithium and silicon, structurally distinct components of the clay lattice, we reveal a multiphasic *in vivo* clearance model for nanoclay biomaterials consistent with progressive remodelling, localised BMP-2 activity and progressive systemic clearance.

## 2. Materials and methods

### 2.1. Materials

### 2.1. Preparation of nanoclay gel

Nanoclay gels were prepared as described previously [5]. Briefly, Laponite^®^ XLG (BYK Widnes, UK) was dispersed in deionized water (dH_2_O) at a concentration of 2.8 wt. % under rapid agitation. The preparations were subsequently sterilized by autoclave, and evaporated water was replaced with sterile dH_2_O.

### 2.2 Preparation and evaluation of labelled BMP-2

#### 2.2.1 BMP-2 dye labelling

Bone morphogenetic protein-2 (BMP-2, carrier-free, Bio-techne, R&D systems, UK) was fluorescently labelled using DyLight™ 800 Amine-reactive Dye (NHS Easter, ThermoFisher Scientific). After dissolving lyophilized BMP-2 in 4 mM HCl, the BMP-2 solution (1 mg/ml) was added to a dialysis cassette (Slide-A-Lyzer™, 10 K MWCO, Thermo Scientific) hydrated in a formulation buffer (0.07% L-glutamic acid, 0.5% sucrose, 0.029% sodium chloride, 0.01% polysorbate).

The cassette was immersed in the formulation buffer and left for 4 hours at 4 °C under stirring to remove excess glycine from the protein and to enhance dye-protein conjugation. Subsequently, the protein was collected from the cassette using a syringe with a needle and mixed with DyLight 800 dissolved in dimethyl sulfoxide (DMSO) in a 5-fold molar excess. The mixture of protein and dye was left on a rotator for 1 hour at room temperature in the dark to induce dye labelling. The reaction mixture was injected into the hydrated dialysis cassette and dialyzed to the formulation buffer for 4 hours at 4 °C under stirring. The labelling degree was determined by fluorophore absorption and the protein absorbance at 280 nm, corrected for the fluorophore using a Nanodrop spectrophotometer (Thermo Scientific). The degree of labelling was defined as the average dye to protein concentration ratio.

#### 2.2.2. Evaluation of dye labelled BMP-2 detection sensitivity

BMP-2 and dye-labelled BMP-2 were subjected to sodium dodecyl sulphate-polyacrylamide gel electrophoresis (SDS-PAGE) and scanned to ensure complete removal of free dye. Briefly, a 12% SDS-polyacrylamide gel was prepared following standard procedures. BMP-2 and dye-labelled BMP-2 were prepared in SDS blue loading buffer with sulfhydryl reagents, dithiothreitol (DTT), and heated at 95 °C for 5 minutes. 50 µl of samples and a protein marker were loaded into each well, and the gel was run at 150 V for 60 minutes (Mini-PROTEAN, Bio-Rad, UK). The gel was fixed in 10% (v/v) acetic acid and 50% methanol for 1 hour and stained with 0.1% Coomassie Brilliant Blue R-250 in 10% acetic acid and 50% methanol for 20 minutes with gentle agitation. After destaining the gel several times, it was scanned. Fluorescence detection was carried out using a Li-Cor Odyssey infrared imager (LI-COR Biosciences Ltd., UK) at the 700 nm channel.

To assess the detection sensitivity of dye-labelled BMP-2 at various concentrations, the labelled BMP-2 (580 µg/ml) was diluted with formulation buffer to prepare concentrations of 290, 145, 72.5, 36.25, 18.13, 9.06, 4.53, 2.27, 1.13, and 0.57 µg/ml in 0.5 ml Eppendorf tubes. All tubes were subsequently scanned using an *in vivo* imaging system (IVIS, Perkin Elmer, Hopkinton, MA) at 740 nm (excitation) and 790 nm (emission) wavelengths. Quantification of dye intensities was performed using Living Image 4.5 software (Perkin Elmer). Fluorescence intensities were quantified within the region of interest (ROI) by calculating the flux radiating omnidirectionally from the ROI in each sample. Calculations were graphically represented as radiant efficiency (photons/s/cm^2^/str)/(μW/cm^2^). The BMP-2 concentration-response curve was interpolated using GraphPad Prism 7 software.

#### 2.2.3. Evaluation of dye labelled BMP-2 activity

The bioactivity of dye-labelled BMP-2 was assessed by the alkaline phosphatase assay. C2C12 cells, mouse muscle myoblasts, were seeded at 5 × 10^4^ cells per well across a 24-well plate and incubated overnight in Dulbecco’s Modified Eagle Medium (DMEM) with 2% fetal calf serum (FCS) and 1% penicillin/streptomycin at 37 °C in a humidified 5% CO_2_ atmosphere. After two washes with PBS, the media was replaced with DMEM containing BMP-2 or labelled BMP-2 dilutions (0, 25, 50, and 100 ng/ml) and cultured for an additional 72 hours (n=3). Cells were fixed in 95% ethanol and stained for alkaline phosphatase, followed by a 60-minute incubation at 37 °C. After stopping the reaction by washing in distilled water, images were captured on a Nikon microscope, and image analysis was conducted using CellProfiler software.

### 2.3 Subcutaneous implantation study of BMP-2 and nanoclay biodistribution

All animal studies were conducted in compliance with the regulations in the Animals (Scientific Procedures) Act 1986 U.K and Home Office license (PPL P96B16FBD) and ethical approval, following institutional and the ARRIVE guidelines. Throughout the studies mice had access to *ad libitum* standard chow and water. Female MF-1 wild-type mice were anesthetized with an intraperitoneal injection of a hypnorm and hypnovel mixture (1:1).

#### 2.3.1 Treatment preparation

Labelled BMP-2 solution at a concentration of 580 μg/ml, was added as a 10% volume fraction to 3.1 wt. % nanoclay gel to give a final concentration of 58 μg/ml BMP-2. A total BMP-2 dose of 2.9 μg was delivered in a 50 μl volume to enable in vivo tracking of the labelled BMP-2. BMP-2 only controls were prepared in the same way substituting saline for the nanoclay gel.

#### 2.3.2 Implantation procedure

Female MF-1 wild-type mice were anesthetized, and subcutaneous injections were performed on the backs of each mouse. Each animal received a single 50 µl implant volume corresponding to a dose of 2.9 μg of labelled BMP-2, with or without nanoclay gel. 4 mice were treated for each time point (4 hours, 1 day, 3 days and 1, 2, 4, 6 and 8 weeks) in the treatment group to allow an *n* of 4 for *in vivo* imaging, an *n* of 3 for ICP-MS and a single sample for histological analysis. In the nanoclay-free control group, where bone induction was not anticipated, the time points of 2, 6, and 8 weeks were excluded from animal numbers in keeping with ARRIVE guidelines.

#### 2.3.3 In vivo imaging of labelled BMP-2

Mice were immediately scanned after sample injection using the *In Vivo* Imaging System (IVIS) at 740 nm (excitation) and 790 nm (emission) wavelengths to capture the initial dye intensity. At each subsequent time point, mice were sacrificed and rescanned to assess dye intensity over time post-injection. Regions of interest (ROI) were defined as the area of samples injected, and the radiant efficiency of the ROI was calculated. The percentage of dye-labelled BMP-2 remaining in the injected area was determined using the following equation:

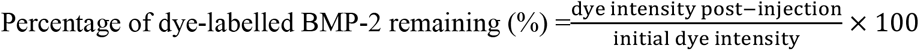

To determine the distribution of labelled BMP-2 in internal organs, including the heart, kidney, liver, lung, spleen, stomach, bladder, and blood, these organs were harvested and collected at each time point. Individual scans were performed, and the radiant efficiency was calculated for each organ.

#### 2.3.4 ICP-MS analysis of nanoclay gel

At each time point (day 1, 3, and weeks 1, 2, 4, 6, and 8), nanoclay gels including the implant site, blood, and organs (heart, kidney, liver, lung, spleen, stomach, bladder) were collected. As a control, tissues, blood, and all organs from female MF-1 wild-type mice without nanoclay gel injection were also collected and used for baseline measures at Day 1 and Week 4.

Harvested organs were frozen, freeze-dried, and weighed (n=3). 2 ml of aqua regia, a nitric acid and hydrochloric acid mixture with a 1:3 ratio, was added to the lyophilized samples. Subsequently, 250 µl of hydrogen peroxide was added, followed by digestion on a hotplate. After cooling, the samples were diluted tenfold with 3% HNO_3_. The Si and Li content were analyzed using Thermo Scientific ELEMENT XR HR-ICP-MS with indium as an internal standard and Phosphorus measured to normalize the amount of biological tissue analyzed.

### 2.4. In vivo bone formation of nanoclay gels

#### 2.4.1 Micro-CT analysis

Nanoclay gels, with and without labelled BMP-2, that were subcutaneously injected were harvested and subjected to micro-computed tomography (micro-CT) analysis using a Skyscan 1176 instrument (Bruker, Kontich, Belgium). The scanning parameters included 45 kV, 556 µA, a 0.2 mm Al filter, and a pixel size of 18 mm. The acquired images were reconstructed using NRecon software, incorporating corrections for misalignment and ring artifacts. For the quantification of bone volume in each sample, the reconstructed images were analyzed using CTAn software. Additionally, CTvox software was employed to generate and visualize 3D models of the samples.

#### 2.4.2 Histology staining

Samples were fixed in 4% paraformaldehyde in PBS at 4 °C for 3 days, dehydrated through a series of ethanol washes (50%, 90%, and 100% in dH_2_O), and incubated in Histo-Clear (National Diagnostics). Following immersion in liquid paraffin at 60 °C, the samples were embedded in a paraffin block for sectioning. Sections were cut at 7 µm using a microtome (Microm 330; Optec, UK).

After rehydration through Histo-Clear and a series of ethanol solutions (100, 90, and 50%), sections were stained for a nuclear counterstain with Weigert’s haematoxylin, followed by staining with 0.5% Alcian Blue 8GX and 1% Sirius Red F3B for proteoglycan-rich matrix and collagenase matrix. Slides were mounted with dibutyl phthalate xylene. Additionally, separate slide sections were stained with Auramine O (Merck) for co-localisation of nanoclay gel within the tissue, followed by mounting with Fluoromount™ (Sigma Aldrich).

#### 2.4.2 Immunofluorescence staining

Slides were deparaffinized and rehydrated, followed by antigen retrieval in heated citrate buffer for 20 minutes (10 mM citrate). After treatment with permeabilization buffer containing 0.3% Triton X-100 in PBS for 5 minutes, slides were blocked in 1% bovine serum albumin in PBS for 5 min. CD68 and CD206 antibodies diluted in 1% BSA in PBS (1:100, Abcam, UK) was added to the slices for pan-macrophage staining. After a 1-hour incubation at room temperature, the slides were washed using PBS and incubated with secondary antibodies—goat anti-rat 488 for CD68 and 647 for CD68 and goat anti-rabbit 647 for CD206. After washing, the nuclei were labelled with DAPI, and the slides were coverslipped with Fluoromount™. All images were captured with a Zeiss Axiovert 100 microscope.

### 2.5 Statistical analysis

All statistical analyses were performed using GraphPad Prism 10. Results are reported as mean ± SD. A p-value < 0.05 was considered statistically significant between different experimental sets, analyzed using one-way and two-way ANOVA with Dunnett’s multiple comparisons test. Systemic biodistribution of lithium and silicon following subcutaneous implantation of a nanoclay–BMP-2 gel, relative to baseline, was assessed by two-way ANOVA with Dunnett’s multiple comparisons test.

## 3. Results and Discussion

To investigate the *in vivo* fate of nanoclay gels and their interaction with BMP-2, we first confirmed the retention of labelled BMP-2, and the induction of bone formation at the implantation site, before examining the clearance kinetics and biodistribution of the nanoclay over the course of the bone remodelling process.

### 3.1. Bone induction through BMP-2 localisation in nanoclay gels

To verify that BMP-2 remains localised within nanoclay gels and induces bone formation *in vivo*, we first tracked dye-labelled BMP-2 over an 8-week period using IVIS imaging and confirmed osteoinductive activity via µCT (**Figure 1**). The amine-reactive near-infrared dye used for BMP-2 labelling was validated for purity, sensitivity (Figure 1a–b), and biological activity using an ALP dose–response assay in C2C12 myoblasts, which confirmed preserved, albeit reduced, activity (Figure 1c). The reduction in BMP-2 bioactivity following amine labelling is likely due to attenuated receptor-binding from lysine modification.

**Figure 1.**
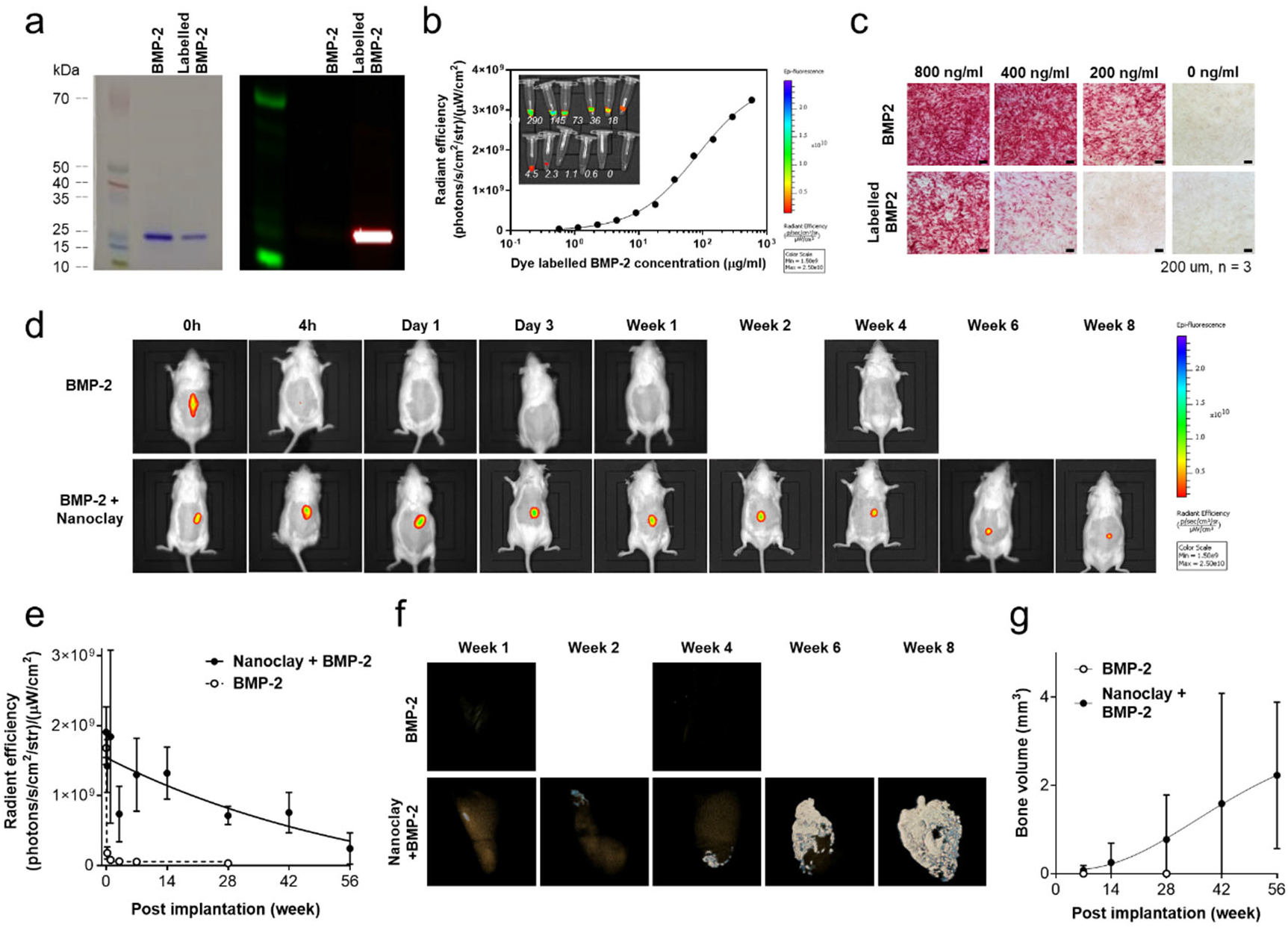
BMP-2 localisation and bone induction in nanoclay gels. (a) SDS-PAGE analysis of native and amine-reactive dye-labelled BMP-2 followed by Coomassie staining and fluorescent gel scanning. Strong bands were observed for both forms upon Coomassie staining, while fluorescence was restricted to the labelled BMP-2. A dilution series of dye-labelled BMP-2 (580–0.57 µg/mL) was imaged using IVIS, confirming a sigmoidal relationship between concentration and radiant efficiency, with a linear response above ∼10 µg/mL and a lower limit of detection (LOD) of ∼4.4 µg/mL (defined as mean baseline signal + 3 SD). (c) Alkaline phosphatase (ALP) assay in C2C12 cells exposed to BMP-2 or labelled BMP-2 (0–800 ng/mL) for 72 h demonstrated retained osteogenic activity of the labelled protein at higher doses. (d) Representative IVIS images and quantification of IVIS radiant efficiency at the injection site over time (e) following subcutaneous injection of labelled BMP-2 alone or with 2.8% nanoclay gel. BMP-2 alone was rapidly lost (undetectable by 4 hours), while BMP-2 with nanoclay remained strongly localised at the injection site up to 8 weeks (n=4). (f) Representative 3D µ-CT reconstructions and bone volume measurements (g) of ectopic bone formed at 8 weeks post-implantation. Robust bone formation was observed in mice receiving BMP-2 with nanoclay gels, while BMP-2 alone showed no detectable bone (n=3).

Subcutaneous injection of labelled BMP-2, with or without nanoclay gels, showed strong fluorescence at the injection site immediately post-injection. However, signal from BMP-2 alone was undetectable by 4 hours, with no signal detectable in blood or organs (**Supplementary Figure S1**), suggesting rapid systemic clearance below lower detection limits (∼4.4 µg/mL). In contrast, BMP-2 co-delivered with nanoclay remained strongly localised at the injection site, with detectable signal persisting even up to 8 weeks (Figure 1d). µ-CT imaging (Figure 1f-g) and histology (**Supplementary Figure 2**) confirmed progressive ectopic bone formation.

### 3.2 Nanoclay degradation and clearance at the implantation site

To define the *in vivo* fate of nanoclay gels following subcutaneous implantation, lithium (Li) and silicon (Si) concentrations at the injection site were quantified over 8 weeks by ICP-MS (**Figure 2**). These two elements represent structurally distinct components of the clay with potentially divergent dissolution profiles: lithium is incorporated within the octahedral sheet of the Laponite lattice, while silicon constitutes the silicate framework (cf. **Figure 5b** below). Tracking these elements provides insight into the temporal sequence of structural degradation of nanoclay gels and their constituent particles against a biological background.

**Figure 2.**
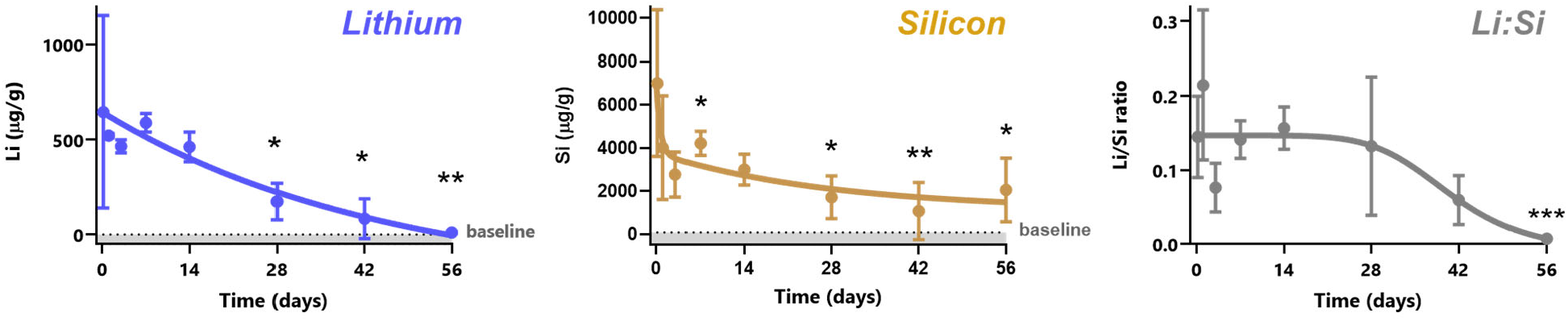
Temporal clearance of lithium and silicon ions from the injection site following subcutaneous implantation of nanoclay-BMP-2 composite gels. ICP-MS analysis was performed on tissue harvested from the injection site over 8 weeks to quantify retained levels of lithium (Li) and silicon (Si), compositional ions of Laponite nanoclay. (a) Lithium levels showed a near-linear decline over time, reaching baseline by 8 weeks. A one-phase exponential decay model (solid line) yielded a decay rate constant (K) of 0.022, and an estimated half-life of 31.47 days (R^2^ = 0.633). Si levels exhibited biphasic clearance, with a two-phase exponential decay model (solid line) indicating a fast component (t½ = 0.45 days) and a slower phase (t½ = 19.6 days), plateauing at ∼1120 µg/g by day 28 (R^2^ = 0.521). The Li:Si mass ratio remained stable during the first 4 weeks, followed by a marked decline. A four-parameter logistic model was fitted to capture the inflection (solid line, IC_50_ ≈ 40.6 days). Statistical comparisons were performed *via* two-way ANOVA with Dunnett’s post hoc test, comparing each timepoint to the 4-hour baseline (* P < 0.05; ** P < 0.01; *** P < 0.001). Data are presented as mean ± SD, N=3 mice per timepoint.

#### 3.2.1 Lithium and silicon follow divergent clearance kinetics from the implant site

Following subcutaneous injection, lithium levels declined progressively and reached baseline by 8 weeks. Fitting a one-phase exponential decay model to the Li data indicated a half-life of 31.47 days (R^2^ = 0.623), indicating steady and continuous clearance. Silicon clearance followed a biphasic pattern with a two-phase exponential decay model providing a better fit (R^2^ = 0.539) than a single-phase model reflecting an initial drop over the first 2-3 days followed by a slower decline approaching a plateau at later time points. The early Si reduction likely reflects the transport of loosely bound or surface-accessible nanoclay particles, while the slower decline in Si indicates longer-term remodelling of the silica-dominant nanoclay gel structure.

Analysis of the Li:Si ratio over time provides further support for this picture, indicating that particles remain stable during the first 4 weeks, before a loss of lithium relative to silicon. This is consistent with clearance *via* transport of intact nanoclay particles (rather than dissolution and diffusion of dissociated ions) during the early weeks post-implantation followed by, at later time-points, some nanoclay particle dissolution at the implant site in which soluble lithium ions are released more efficiently than silicon. The latter may persist as particulate remnants or polymerise and form less mobile amorphous silica species and may, in addition, be likely to integrate with newly formed bone tissue. The, perhaps surprising, lack of appreciable nanoclay dissolution at the implantation site itself over the first four weeks is consistent with the known concentration-dependent chemical stability of Laponite, where higher nanoclay concentrations (>2 wt%) and ionic environments suppress proton-driven degradation of the crystalline lattice [23, 24].

#### 3.2.2 Histological visualisation of nanoclay gel persistence and clearance

Histological analysis using Auramine O staining, in parallel with H&E, A&S and CD68 immunohistochemistry, was used to assess clearance and remodelling of the nanoclay gel over the course of BMP-2-induced bone formation (**Figure 3 and 4**). Auramine O is a cationic fluorochrome with strong affinity for clay surfaces [25, 26]. When staining histological sections, omitting the acid–alcohol differentiation step used classically for Auramine O staining of acid-fast bacteria, results in retention of Auramine O within both nanoclay and surrounding tissue, but with markedly distinct emission characteristics. Under blue excitation with long-pass detection, nanoclay-rich regions exhibit intense orange fluorescence, enabling high-contrast visualisation of extra- and intracellular nanoclay aggregates against a blue–green tissue background [27].

**Figure 3.**
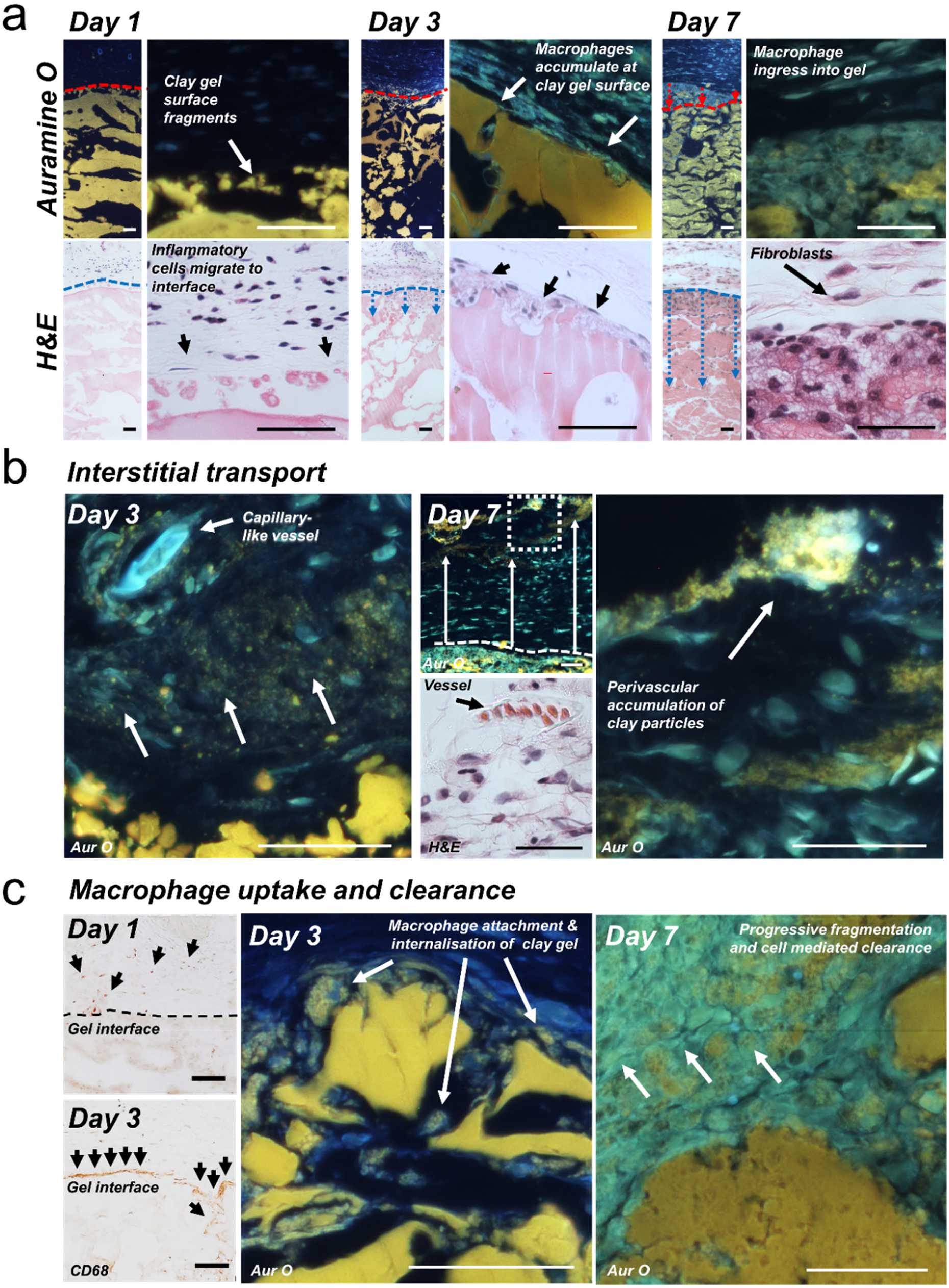
Cellular invasion, fragmentation, and macrophage-mediated clearance of the nanoclay gel implant. Representative Auramine O (Aur O)–stained and H&E-stained histological sections illustrating early interactions between host tissue and the nanoclay gel at days 1, 3 and 7 post injection. (**a**) Low- and higher-power images at the nanoclay gel–tissue interface show progressive fragmentation and cellular remodelling. Annotations indicate progressing boundary of the gel (red) and cell infiltrate (blue). (**b**) Interstitial transport of nanoclay fragments accumulating around capillary-like vascular structures. (**c**) Macrophage (CD68-positive cells) recruitment and uptake of nanoclay and cell-mediated clearance. Scale bars = 100μm.

**Figure 4.**
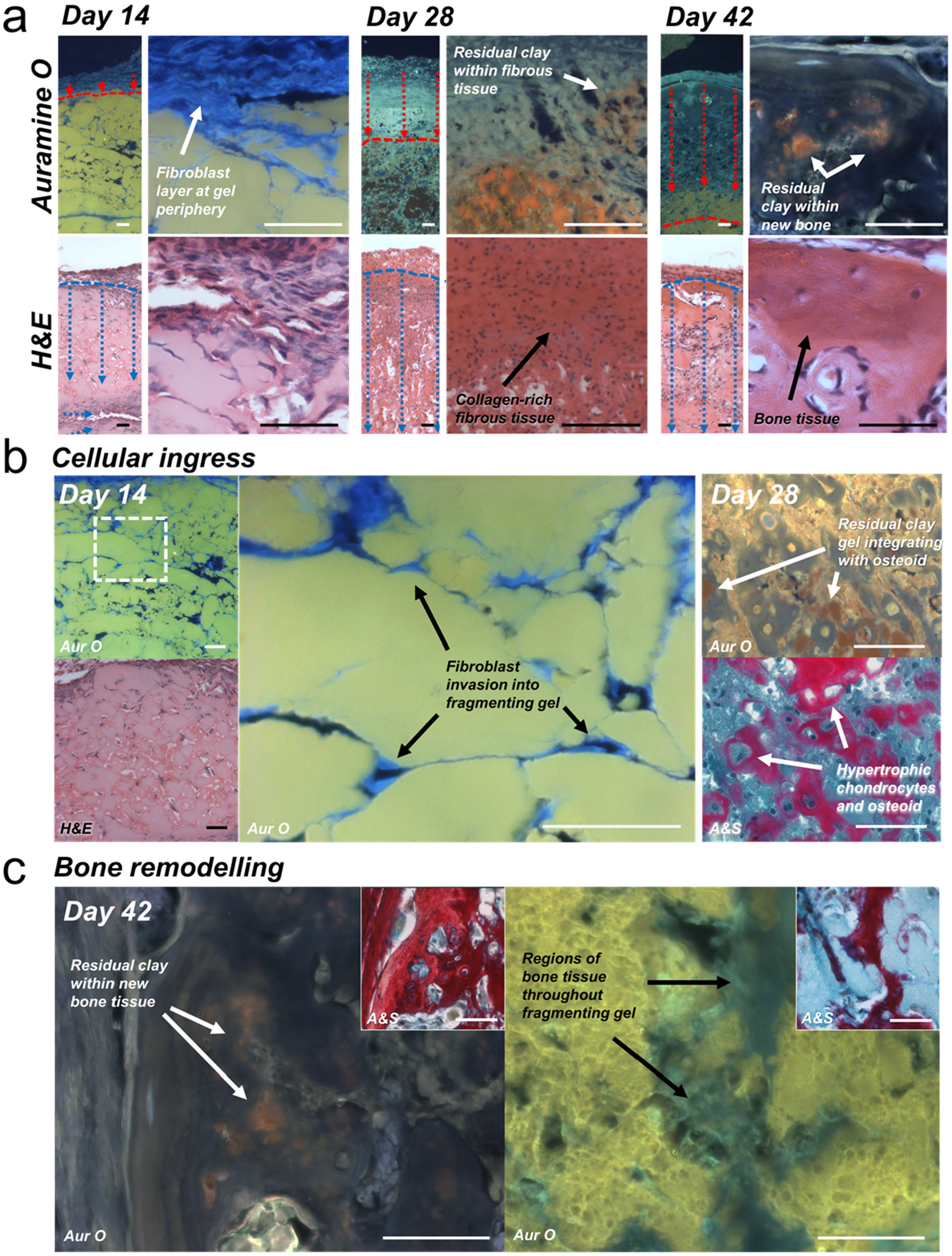
Fibroblast-mediated remodelling and bone formation within the nanoclay gel implant. Representative Auramine O (Aur O) stained, H&E-stained, and Alcian blue Sirius red (A&S) stained histological sections illustrating remodelling of the nanoclay gel at days 14, 28, and 42 post injection. (**a**) Low and higher-power images at the gel-tissue interface show progressive cellular invasion and remodelling of the nanoclay gel. Annotations indicate the gel boundary (red) and infiltrating tissue (blue). (**b**) Fibroblast colonisation of the fragmenting nanoclay gel, showing a dense fibroblast layer at the gel periphery and fibroblast penetration between gel fragments. (**c**) Bone formation and remodelling, with residual nanoclay integrated within, or contiguous with, newly formed bone. Scale bars = 100 μm.

#### 3.2.3 Early macrophage recruitment and progressive fragmentation of the nanoclay gel

Histological analysis of early timepoints indicate the nanoclay gel undergoes progressive, cell-mediated clearance and remodelling over the first week following implantation (**Figure 3**). At Day 1, Auramine O staining showed an intact but irregular gel surface, with discrete fragments apparent at the interface between the bulk gel and surrounding tissue. H&E demonstrated infiltration of acute inflammatory cells migrating toward the gel surface. CD68 immunostaining confirmed the presence of macrophages adjacent to, but not yet extensively interacting with the gel, indicating early recruitment to the implant site (Figure 3a & c). By Day 3, Auramine O and H&E staining showed macrophages accumulating along the gel surface, forming a contiguous cellular layer and beginning to penetrate into disrupted regions of the gel. CD68-positive cells densely lined the interface, and higher-power Auramine O images demonstrated close apposition of macrophages to fragmented nanoclay gel structures. Auramine O-positive material was present within vacuolated macrophages indicating active phagocytosis of nanoclay. By Day 7, macrophage ingress into the gel was more extensive. Auramine O staining revealed loss of a continuous gel boundary, with infiltration of cells between larger nanoclay gel fragments and accumulation of Auramine O-positive material within densely packed cellular regions. Numerous granulated Auramine O positive cells were observed adjacent to, and removed from, the bulk nanoclay gel, consistent with progressive fragmentation and cell-mediated clearance of nanoclay gel.

In addition to local clearance at the gel interface, evidence of interstitial transport of nanoclay fragments was also observed. At Day 3, fine punctate Auramine O positive material could be detected between cells and in regions proximal to the gel, frequently localising around nearby capillary-like vascular structures adjacent to the gel–tissue interface. By Day 7, larger accumulations were present at sites distant from the gel-tissue interface, with pronounced perivascular association. This spatial redistribution suggests active transport of fragmented nanoclay away from the implant site, likely mediated by infiltrating macrophages.

#### 3.2.4 Transition to fibroblast-driven remodelling and bone formation

At later time points, histological analysis revealed a marked transition from the early inflammatory and macrophage-mediated clearance phase to fibroblast-driven remodelling and subsequent bone formation within the nanoclay gel implant (**Figure 4**). By Day 14, accompanying ongoing macrophage activity, fibroblast accumulation at the interface was observed with extensive penetration into the fragmenting gel structure. Within the nanoclay gel, individual cells exhibited elongated morphologies with long cytoplasmic processes, forming interconnected networks spanning between gel fragments. The extent of integration and direct cell-gel interaction suggests a fibroblastic remodelling rather than foreign body encapsulation response to the nanoclay gel. At Day 28, further remodelling was evident, with regions of osteochondral tissue closely integrated with residual nanoclay. Auramine O-positive gel fragments were observed in direct apposition to osteoid-like matrix and hypertrophic chondrocyte-like cells identified in H&E sections. By Day 42, extensive bone tissue formation was observed throughout the former nanoclay gel volume. Alcian blue Sirius red staining confirmed the presence of collagen-rich, mineralised bone tissue, while Auramine O staining revealed residual nanoclay incorporated within newly formed bone as well as regions where bone formed contiguous with remaining gel fragments. The co-existence of bone tissue and Aur O–positive material suggests that persistent silica structures become structurally integrated within the early woven bone, rather than being completely excluded.

#### 3.2.5 Multiphasic nanoclay clearance and growth-factor presentation in vivo

Taken together, the ICP-MS and histological analysis support a picture of progressive, multiphasic clearance from the implant site. The early drop in silicon and lithium detected by ICP-MS coincides with histological evidence of early gel fragmentation at the surface and clearance of nanoclay particulates via interstitial transport over the first few days. This is followed by a more sustained period of macrophage phagocytic activity driving, with fibroblastic invasion, progressive fragmentation of the gel. The stable Li:Si ratio together with observations of macrophages transporting Auramine O-positive material implies clearance *via* uptake of disaggregated particulates rather than ionic dissolution of the clay platelets themselves during this phase (**Figure 5a**). The later decoupling of the Li:Si ratio at around 28 days thus indicates a further transition in which local nanoclay dissolution becomes a factor, with Li^+^ ions displaying more efficient release than silicon from the implant site (**Figure 5b**). Interestingly, this transition coincides with the nanoclay gel becoming increasingly embedded within new osteoid-rich matrix possibly accounting for the slower release of silicon through entrapment of less mobile silica species, or - as suggested by residual Auramine O staining - through incorporation into the newly formed bone matrix itself.

**Figure 5.**
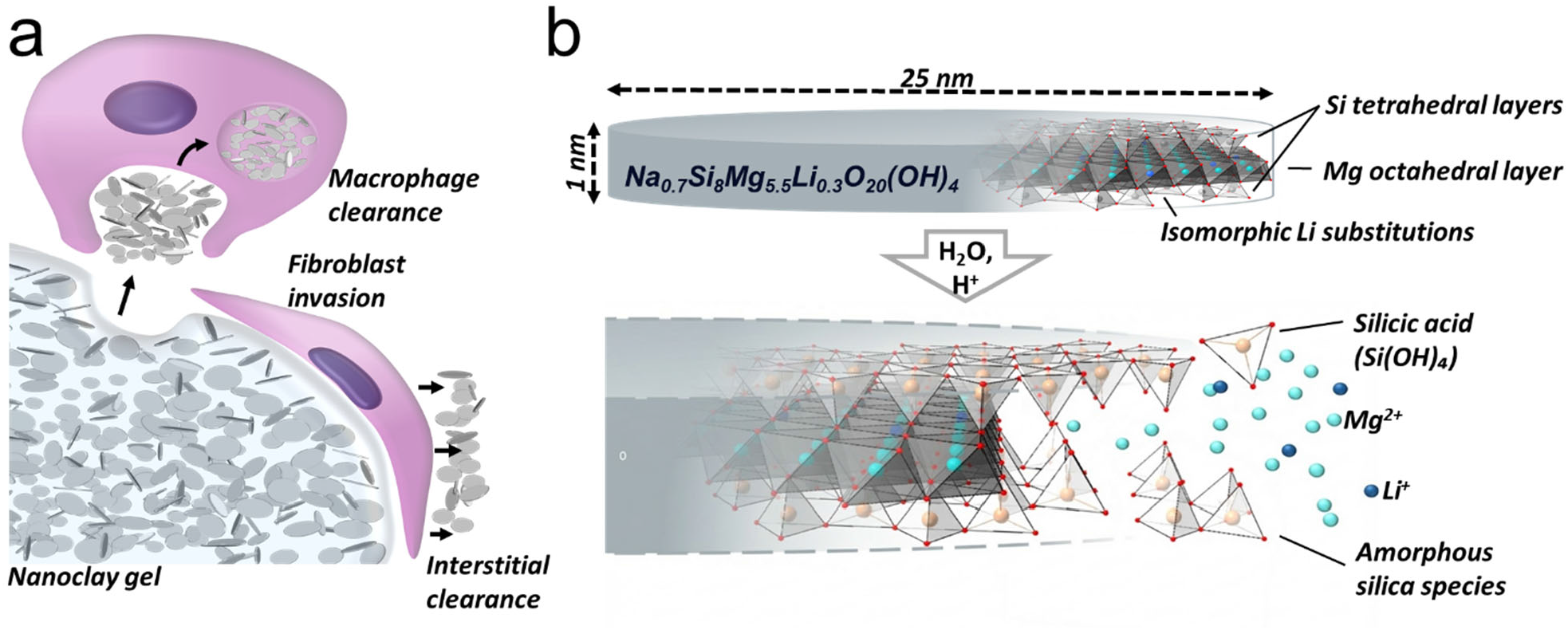
Mechanisms of nanoclay gel clearance. Following implantation, nanoclay gel fragmentation and clearance occurs interstitially and through direct interaction with infiltrating cells. Macrophages internalise and transport nanoclay particulates and fibroblasts progressively penetrate and span the fragmenting gel structure (**a**). Over the course of clearance, nanoclay platelets undergo dissolution resulting in the release of lithium ions alongside silica-containing species yielding divergent biodistribution profiles (**b**).

This analysis also provides some clues as to how nanoclay-sequestered BMP-2 becomes accessible to cells despite limited release from the gel. Histology demonstrates close interactions between the fragmenting gel and infiltrating macrophages, fibroblasts, and osteogenic cells, suggesting that cellular attachment, phagocytic processing, and matrix remodelling progressively expose nanoclay-bound BMP-2 at the cell-material interface. This interpretation is consistent with *in vivo* imaging data showing prolonged localisation of labelled BMP-2 alongside histological evidence of sustained osteoinductive activity throughout the gel volume. Together, these findings support a model in which the nanoclay gel templates new bone by presenting BMP-2 locally *via* direct cell-nanoclay gel interactions and progressive gel remodelling while limiting BMP-2 release beyond the boundary of the implant.

### 3.3 Nanoclay biodistribution

Alongside analysis of clearance from the implant site, the systemic biodistribution of lithium and silicon was quantified *via* ICP-MS across major organs over 8 weeks (**Figure 6**). Notably, and in contrast to the initial stable Li:Si ratios observed above, divergent biodistribution kinetics for lithium and silicon were observed systemically as early as 4 h post-injection, indicating rapid decoupling of nanoclay dissolution products over the course of clearance and transport.

**Figure 6.**
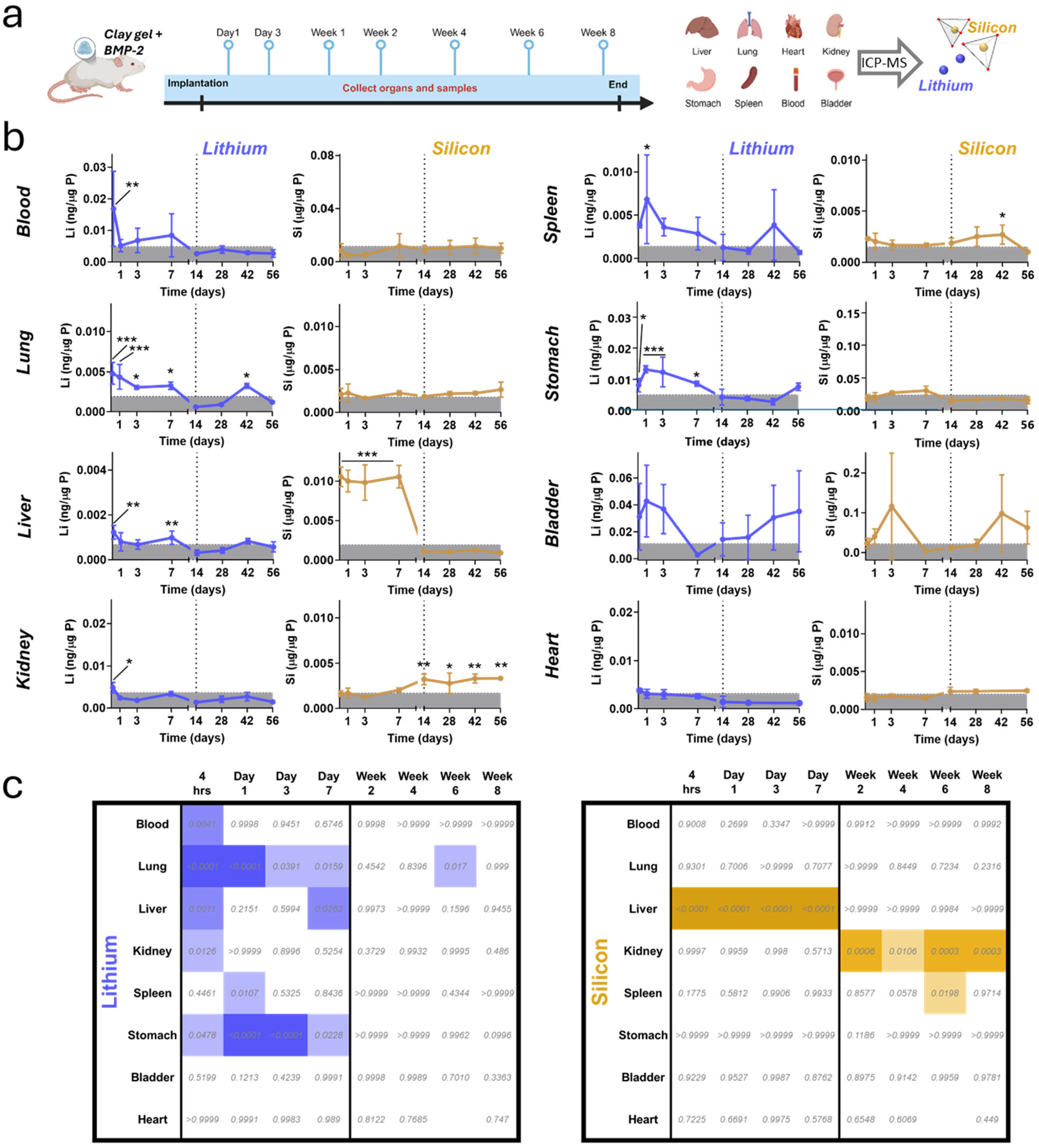
Systemic biodistribution of lithium and silicon following subcutaneous implantation of a nanoclay–BMP-2 gel. Lithium exhibits early, transient systemic distribution consistent with rapid release and clearance following structural degradation of the nanoclay, while silicon shows initial hepatic and subsequent renal involvement consistent with progressive breakdown and handling of the silicate framework. **(a)** Schematic overview of the implantation and sampling timeline. (**b**) Lithium and Silicon concentrations were quantified by ICP-MS and nomalised to Phosphorus content in blood, lung, liver, kidney, spleen, stomach, bladder and heart at 4 h, 1 day, 3 days, 7 days, 14 days, 28 days, 42 days, and 56 days post-implantation. Data are presented as mean ± SD and compared to baseline tissue levels (mean values at 4hrs and day 28) over time (n = 3 per time point). For visualisation days 1–9 and days 14–56 are presented as separate intervals, each scaled to occupy 50% of the x-axis. Grey shaded regions indicate physiological background ranges (mean + SD). Statistical significance relative to baseline was assessed by two-way ANOVA with Dunnett’s multiple comparisons test. P values indicating significant readings above the baseline are indicated * P < 0.05; ** P < 0.01; *** P < 0.001 and summarised in the heatmap (**c**).

Lithium exhibited an early and broadly distributed systemic profile consistent with rapid release into the circulation. By 4 h, Li was significantly elevated above baseline in the blood, lung, liver, kidney and stomach (albeit orders of magnitude below pharmacological dosing levels). Over the next seven days, elevated Li was detected in the lung, liver, spleen, and stomach after which, by day 14, levels were indistinguishable from baseline except for a brief signal in the lung (day 42). In contrast, the early biodistribution of silicon was more specifically defined with a pronounced elevation in the liver, and only the liver, during the first week (4 h, 1 day, 3 days, and 7 days). A marked transition was then observed in the latter phase with detection in the kidney across the time frame (weeks 2, 4, 6 and 8) and transient detection in the spleen (day 42). Elevated means for both silicon and lithium were observed in the bladder at various early and later time points but with large standard deviations, likely due to differences in urine content at the time of sampling.

Lithium and silicon are both, along with magnesium, the structural components of the Laponite crystalline lattice and so the decoupling of their biodistribution profiles indicates, at least some, nanoclay dissolution from the earliest time points. The persistence of stable Li:Si ratios locally and observations of macrophage-driven particulate clearance indicates that Laponite particulates are transported away from the implant before significant dissolution of the particles themselves occurs intracellularly. This interpretation is supported by recent *in vitro* experiments demonstrating that Laponite nanoparticles are internalised as aggregates by cells and processed within endo-lysosomal compartments, where both a reduction in particle size and falling Mg:Si ratio observed *via* TEM-EDX, support intracellular degradation [20].

The chemical basis for this behaviour is well established. Classical studies of nanoclay dissolution and ion transport across dialysis membranes [28] demonstrate that Laponite undergoes dissolution at pH < 9, with proton attack initiating at platelet edges to release the soluble structural cations (**Figure 5b**). Silicon displays retarded mobility across membranes in these models due to the slower breakdown of the tetrahedral layer and, possibly, the reprecipitation of polymeric amorphous silicas. This correlates well with the redistribution profiles observed in the current study where lithium solubilises quickly and enters circulation, whereas early liver-dominant silicon indicates hepatic sequestration of particulate or polymeric silica. At later stages, further solubilisation of silica towards, ultimately, orthosilicic acid, enables subsequent renal handling [29]. The persistence of detectable silicon in the kidney at the terminal time point, is consistent with ongoing renal clearance of residual nanoclay from the implantation site, indicating that slow, ongoing degradation and release of silicate species continues beyond 8 weeks in this model.

Together, these observations support a multistage clearance model in which nanoclay gels are first stabilised locally, then transported and degraded intracellularly, producing chemically distinct degradation products whose divergent biodistribution profiles reflect different solubility, mobility, and clearance pathways.

## 4. Conclusion

This study defines the *in vivo* fate of a synthetic nanoclay–BMP-2 gel during bone induction and demonstrates that nanoclay clearance proceeds through multiphasic, component-decoupled pathways. Early cell-mediated fragmentation and transport of clay particulates dominate clearance from the implant site, while later stages involve preferential lithium release associated with local nanoclay dissolution and prolonged persistence of silica-containing species. These chemically distinct degradation products exhibit divergent systemic handling, with rapid, transient renal clearance of lithium contrasted by initial hepatic and then later phase renal processing of silicon species. Importantly we found no evidence of any sustained off-target accumulation. Together, these findings show how progressive nanoclay remodelling can support stable, localised BMP-2 activity while enabling controlled clearance and safe systemic handling. The results highlight the importance of component-resolved analysis when evaluating the *in vivo* fate and safety of degradable inorganic biomaterials.

## Credit authorship contribution statement

**Yang-Hee Kim**: Conceptualization, Data curation, Formal analysis, Investigation, Methodology, Writing - original draft, Writing - review & editing. **Juan Aviles Milan**: Data curation, Formal analysis, Investigation. **Janos Kanczler**: Data curation, Formal analysis, Investigation, Writing - review & editing. **Agnieszka Janeczek**: Conceptualization, Project administration, Writing - original draft, Writing - review & editing. **Richard O.C. Oreffo**: Project administration, Writing - original draft, Writing - review & editing. **Jonathan I. Dawson**: Conceptualization, Data curation, Formal analysis, Investigation, Methodology, Project administration, Writing - original draft, Writing - review & editing.

## Declaration of competing interest

A.A Janeczek, R.O.C. Oreffo and J.I Dawson are co-founders and shareholders in a University spin out company, Renovos Biologics Ltd. with a license to IP indirectly related to the current manuscript.

## Declaration of AI and AI-assisted technologies in the writing process

During the preparation of this work the author(s) used CHATGPT version 5.3 in order to improve the readability and clarity of the text and to generate graphical components of the graphical abstract. In all cases, the authors reviewed and edited any generated content and take full responsibility for the final content of the publication.

## Acknowledgements

This study was supported by grants from the Biotechnology and Biological Sciences Research Council (BBSRC LO21071/ and BB/L00609X/1) and UK Regenerative Medicine Platform Hub Acellular Approaches for Therapeutic Delivery (MR/K026682/1) Acellular Hub, SMART materials 3D architecture (MR/R015651/1) and the UK Regenerative Medicine Platform (MR/L012626/1 Southampton Imaging) to ROCO and, MRC-AMED Regenerative Medicine and Stem Cell Research Initiative (MR/ V00543X/1) to JID, ROCO and YK. YK acknowledges supports by an Innovate UK grant to Renovos (Innovation in Health and Life Sciences, Project number 103349).

## Supplementary figures

**Supplementary Figure S1.**
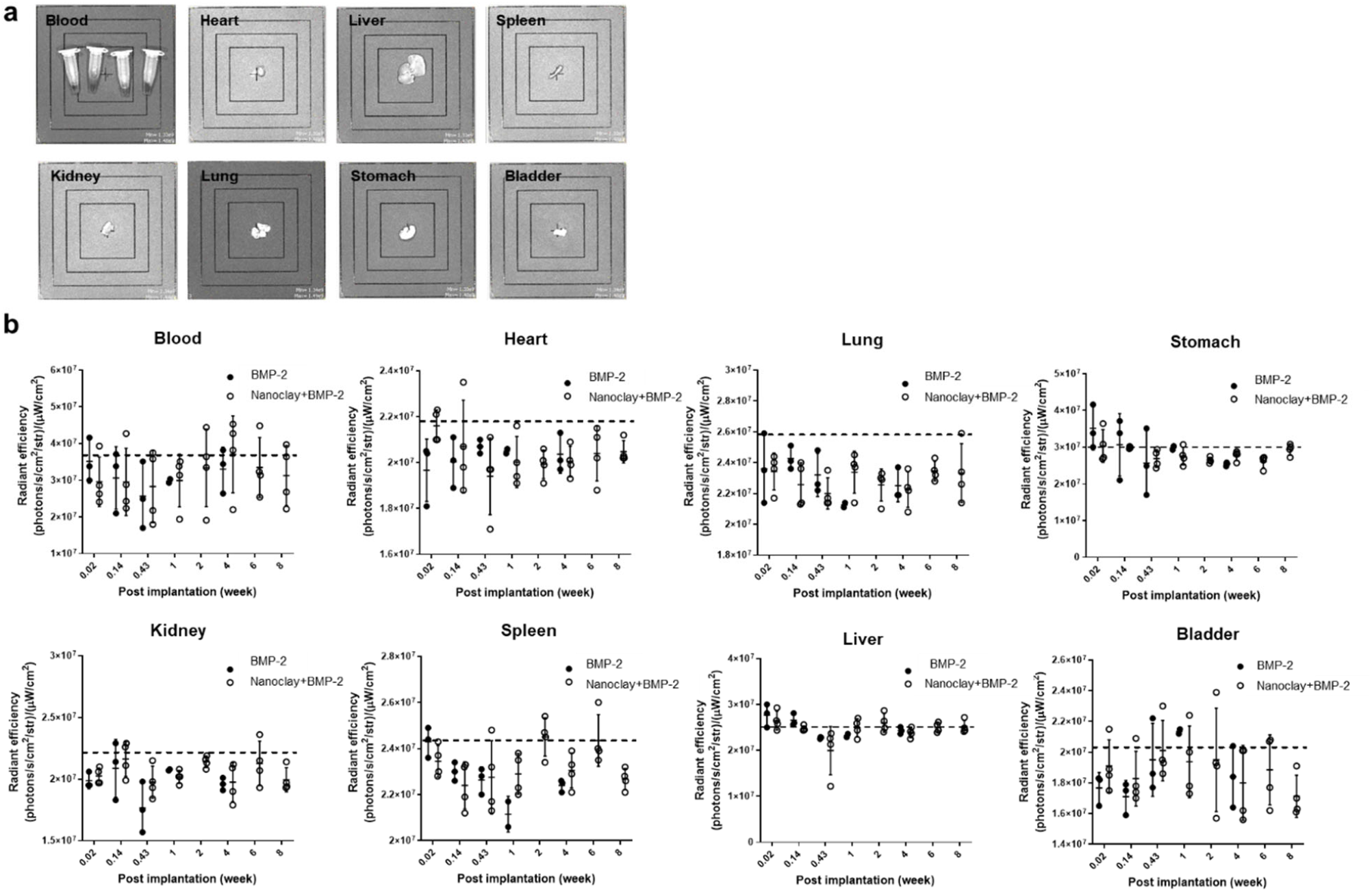
*In vivo* biodistribution of labelled BMP-2 in internal organs. (a) After injection of labelled BMP-2 and 2.8% nanoclay gels with labelled BMP-2, mice were sacrificed and organs (heart, kidney, liver, lung, spleen, stomach, bladder) and blood were harvested and collected at each timepoint, followed by IVIS scanning. No dye signal detected was observed. (b) Radiant efficiencies of ROI were calculated and a dashed line indicates a radiant efficiency from control group without any sample. There is no difference between control group and experiment groups.

**Supplementary Figure S2.**
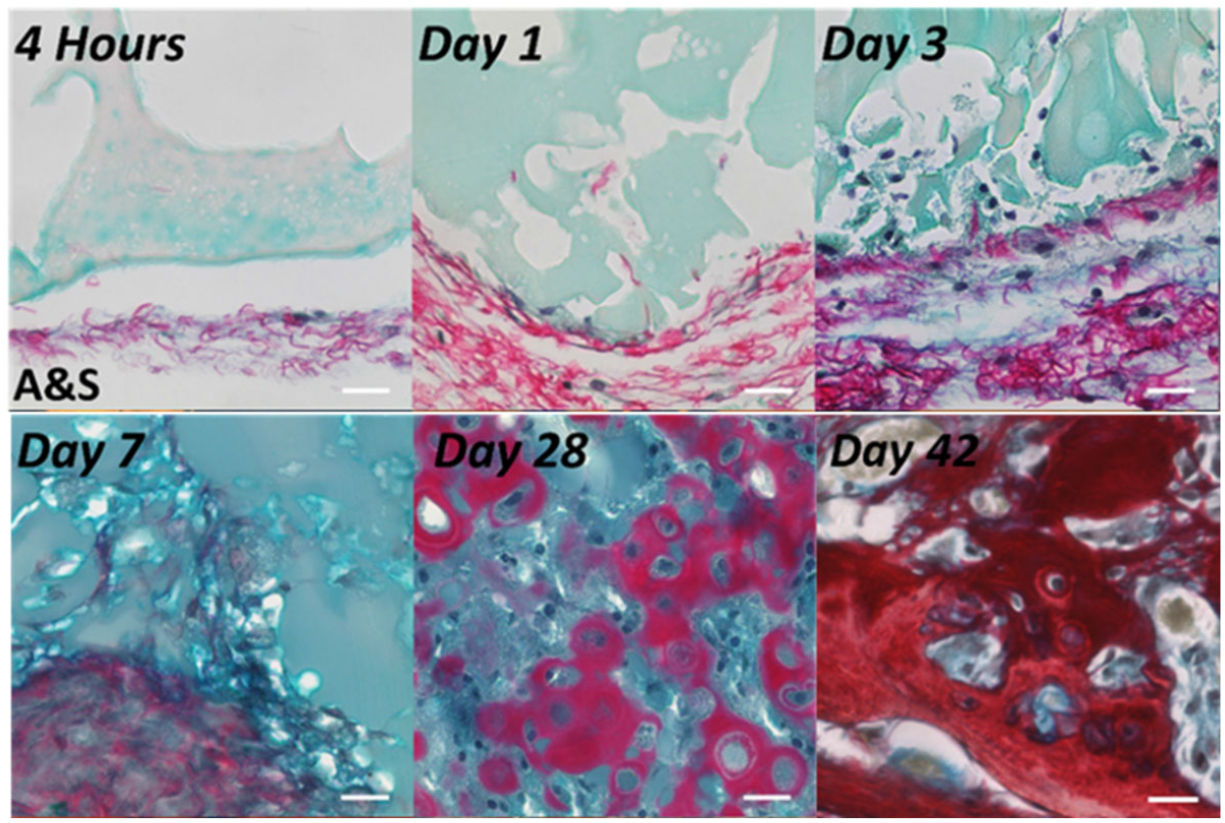
Alcian Blue and Sirius Red (A&S) staining confirms confirmed progressive ectopic bone formation over 42 days following subcutaneous injection of labelled BMP-2 in nanoclay gels.

